# Heterogeneous synaptic weighting improves neural coding in the presence of common noise

**DOI:** 10.1101/811364

**Authors:** Pratik S. Sachdeva, Jesse A. Livezey, Michael R. DeWeese

## Abstract

Simultaneous recordings from the cortex have revealed that neural activity is highly variable, and that some variability is shared across neurons in a population. Further experimental work has demonstrated that the shared component of a neuronal population’s variability is typically comparable to or larger than its private component. Meanwhile, an abundance of theoretical work has assessed the impact shared variability has upon a population code. For example, shared input noise is understood to have a detrimental impact on a neural population’s coding fidelity. However, other contributions to variability, such as common noise, can also play a role in shaping correlated variability. We present a network of linear-nonlinear neurons in which we introduce a common noise input to model, for instance, variability resulting from upstream action potentials that are irrelevant for the task at hand. We show that by applying a heterogeneous set of synaptic weights to the neural inputs carrying the common noise, the network can improve its coding ability as measured by both Fisher information and Shannon mutual information, even in cases where this results in amplification of the common noise. With a broad and heterogeneous distribution of synaptic weights, a population of neurons can remove the harmful effects imposed by afferents that are uninformative about a stimulus. We demonstrate that some nonlinear networks benefit from weight diversification up to a certain population size, above which the drawbacks from amplified noise dominate over the benefits of diversification. We further characterize these benefits in terms of the relative strength of shared and private variability sources. Finally, we studied the asymptotic behavior of the mutual information and Fisher information analytically in our various networks as a function of population size. We find some surprising qualitative changes in the asymptotic behavior as we make seemingly minor changes in the synaptic weight distributions.

## 1 Introduction

Variability is a prominent feature of many neural systems – neural responses to repeated presentations of the same external stimulus will typically vary from trial to trial [41]. Furthermore, neural variability often exhibits pairwise correlations, so that pairs of neurons are more (or less) likely to be co-active than they would be by chance if their fluctuations in activity to a repeated stimulus were independent. These so-called “noise correlations” (which we also refer to as “shared variability”) have been observed throughout the cortex [4, 13], and their presence has important implications for neural coding [1, 52].

If the activities of individual neurons are driven by a stimulus shared by all neurons but corrupted by noise that is independent for each neuron (so-called “private variability”), then the signal can be recovered by simply averaging the activity across the population [1, 32]. If instead some variability is shared across neurons (*i.e.*, there are noise correlations), naively averaging the activity across the population will not necessarily recover the signal, no matter how large the population [52]. An abundance of theoretical work has explored how shared variability can be either beneficial or detrimental to the fidelity of a population code (relative to the null model of only private variability amongst the neurons), depending on its structure and relationship with the tuning properties of the neural population [1, 5, 12, 14, 17, 34, 44, 51, 52].

One general conclusion of this work highlights the importance of the geometric relationship between noise correlations and a neural population’s signal correlations [4, 22]. To illustrate this, the mean responses of a neural population across a variety of stimuli (*i.e.*, those responses represented by receptive fields or tuning curves) can be examined in the neural space (Fig. 1a, black curves). The correlations amongst the mean responses for different stimuli specify the signal correlations for a neural population [4]. Private variability exhibits no correlational structure, and thus its relationship with the signal correlations is determined by the mean neural activity and the individual variances (Fig. 1a, left). Shared variability, however, may reshape neural activity to lie, for example, orthogonal to the mean response curve (Fig. 1a, middle). In the case of Figure 1a, middle, neural coding is improved (relative to private variability), because the variability occupies regions of the neural space that are not traversed by the mean response curve [33]. Shared variability can also harm performance, however. Recent work has identified *differential correlations* – those that are proportional to the products of the derivatives of tuning functions (Fig. 1a, right) – as particularly harmful to the performance of a population code [34]. While differential correlations are consequential, they may serve as a small contribution to a population’s total shared variability, leaving “non-differential correlations” as the dominant component of shared variability. [27].

**Figure 1:**
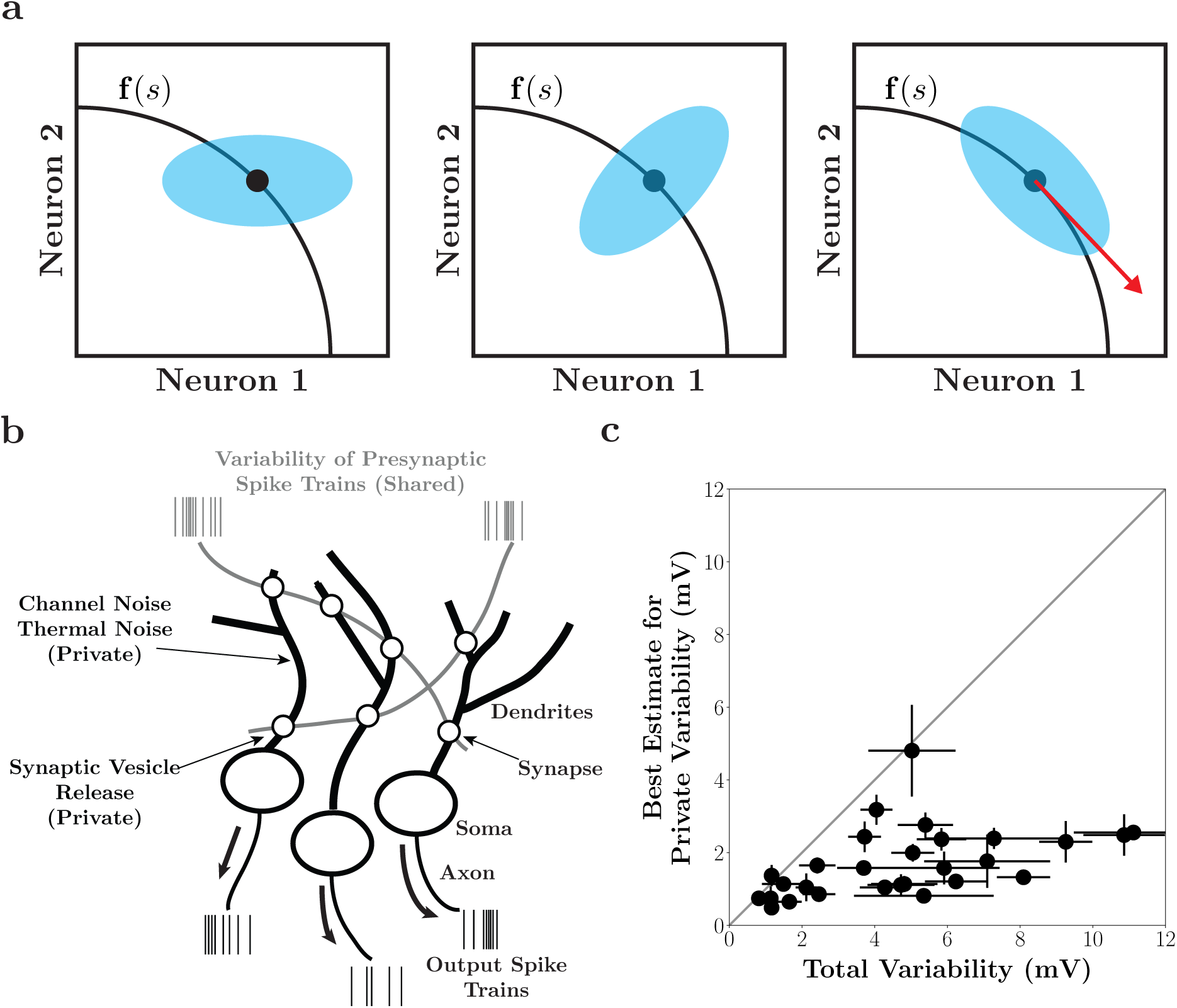
Private and shared variability. **(a)** The geometric relationship between neural activity and shared variability. Black curves denote mean responses to different stimuli. Variability for a specific stimulus (black dot) may be private (left), shared (middle), or take on the structure of differential correlations (right). The red arrow represents the tangent direction of the mean stimulus response. **(b)** Schematic of the types of variability that a neural population can encounter. The variability of a neural population contains both private components (*e.g.*, synaptic vesicle release, channel noise, thermal noise, etc.) and shared components (*e.g.*, variability of pre-synaptic spike trains, shared input noise). Shared variability can be induced by the variability of afferent connections (which is shared across a postsynaptic population) or inherited from the stimulus itself. Furthermore, shared variability is shaped by synaptic weighting. **(c)** Estimates of the private variability contributions to the total variability of neurons (*N* = 28) recorded from auditory cortex of anesthetized rats. Diagonal line indicates the identity. Figure reproduced from [16].

The sources of neural variability – and their respective contributions to the private and shared components – will have a significant impact on shaping the geometry of the population’s correlational structure, and therefore its coding ability [10]. For example, private sources of variability such as channel noise or stochastic synaptic vesicle release could be averaged out by a downstream neuron receiving input from the population [18]. However, sources of variability shared across neurons – such as the variability of presynaptic spike trains from neurons that synapse onto multiple neurons – would introduce shared variability and place different constraints on a neural code [24, 41]. In particular, differential correlations are typically induced by shared input noise (*i.e.*, noise carried by a stimulus) or suboptimal computations [7, 24].

Past work has examined the contributions of private and shared sources to variability in cortex [2, 16]. Specifically, by partitioning sub-threshold variability of a neural population into private components (synaptic, thermal, channel noise in the dendrites, and other local sources of variability) and shared components (variability induced by afferent connections), it was found that the private component of the total variability was quite small, while the shared component can be much larger (Fig. 1b and c). Thus, neural populations must contend with the large shared component of a neuron’s variability. The incoming structure of shared variability and its subsequent shaping by the computation of a neural population is an important consideration for evaluating the strength of a neural code [54].

Moreno-Bote et al. demonstrated that shared input noise is detrimental to the fidelity of a population code [34]. Here, we instead examine sources of shared variability which do not necessarily result in differential correlations (*i.e.*, they do not appear as shared input noise) and thus can be manipulated by features of neural computation such as synaptic weighting. We refer to these noise sources as “common noise” to distinguish them from the aforementioned special case of “shared input noise” [29, 46]. For example, a common noise source could include an upstream neuron whose action potentials are “noisy” in the sense that they are unimportant for the computation of the current stimulus. Common noise, because it is manipulated by synaptic weighting, can serve as a source of nondifferential correlations (*e.g.*, Fig. 1a, middle), thereby having either a beneficial or harmful impact on the strength of the population code. We aim to better elucidate the nature of this impact.

We consider a linear-nonlinear architecture [25, 36, 37] and explore how its neural representation is impacted by both a common source of variability and private noise sources affecting individual neurons independently. This simple architecture allowed us to analytically assess coding ability using both Fisher information [1, 48, 49, 51], and Shannon mutual information. We evaluated the coding fidelity of both the linear representation and the nonlinear representation after a quadratic nonlinearity as a function of the distribution of synaptic weights that shape the shared variability within the representations [35]. We find that the linear stage representation’s coding fidelity improves with diverse synaptic weighting, even if the weighting amplifies the common noise in the neural circuit. Meanwhile, the nonlinear stage representation also benefits from diverse synaptic weighting in a regime where common noise may be amplified, but not too strongly. Moreover, we found that the distribution of synaptic weights that optimized the network’s performance depended strongly on the relative amount of private and shared variability. In particular, the neural circuit’s coding fidelity benefits from diverse synaptic weighting when shared variability is the dominant contribution to the variability. Together, our results highlight the importance of diverse synaptic weighting when a neural circuit must contend with sources of common noise.

## 2 Methods

All code used for the analyses described in this paper is publicly available on Github.^1^

### 2.1 Network Architecture

We consider the linear-nonlinear architecture depicted in Figure 2. The inputs to the network consist of a stimulus *s* along with common (Gaussian) noise *ξ*_*C*_. The *N* neurons in the network take a linear combination of the inputs and are further corrupted by i.i.d. private Gaussian noise. Thus, the output of the linear stage for the *i*th neuron is

**Figure 2:**
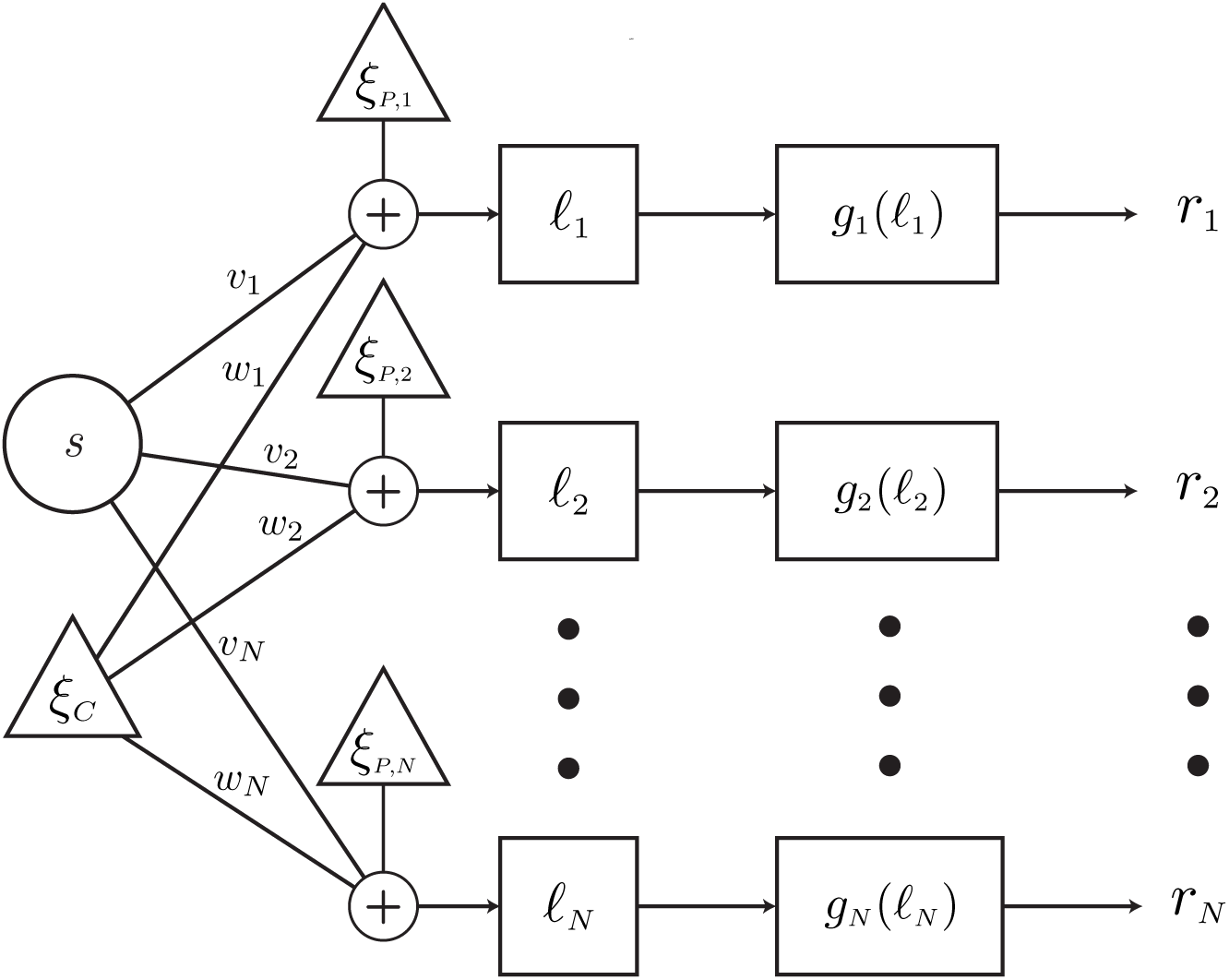
Linear-nonlinear Network Architecture. The network takes as its inputs a stimulus *s* and common noise *ξ*_*C*_. A linear combination of these quantities is corrupted by individual private noises *ξ*_*P,i*_. The output of this linear stage is then passed through a nonlinearity *g*_*i*_(ℓ) to produce a “firing rate” *r*_*i*_. The weights for the linear stage of the network, *v*_*i*_ and *w*_*i*_, can be thought of as synaptic weighting. Importantly, the common noise is distinct from shared input noise because it is manipulated by the synaptic weighting.

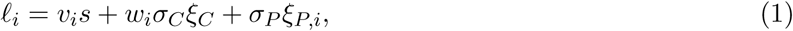

where *ξ*_*P,i*_ is the private noise, *v*_*i*_ and *w*_*i*_ are the weights, and the common and private noise terms are scaled by positive constants *σ*_*C*_ and *σ*_*P*_. The noisy linear combination is passed through a nonlinearity *g*_*i*_(ℓ_*i*_) whose output *r*_*i*_ can be thought of as a firing rate.

Thus, the network-wide computation is given by

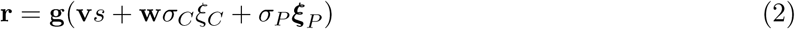

where **g**(ℓ) is an element-wise application of the network nonlinearity.

### 2.2 Measures of Coding Strength

In order to assess the fidelity of the population code represented by ℓ or **r**, we turn to the Fisher information and the Shannon mutual information [15]. The former has largely been utilized in the context of sensory decoding and correlated variability [1, 4, 27] while the latter has been well studied in the context of efficient coding [3, 6, 9, 39].

The Fisher information sets a limit by which the readout of a population code can determine the value of the stimulus. Formally, it sets a lower bound to the variance of an unbiased estimator for the stimulus. In terms of the network architecture, the Fisher information of the representation **r** (or ℓ) quantifies how well *s* can be decoded given the representation. For Gaussian noise models with slowly varying covariances, the Fisher information is equal to the linear Fisher information (LFI):

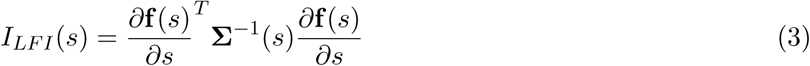

where **f** (*s*) and **Σ**(*s*) are the mean and covariance of the response (here **r** or ℓ) to the stimulus *s*. In other cases, the LFI serves as a lower bound for the Fisher information and thus is a useful proxy when the Fisher information is challenging to calculate analytically. The estimator for *I*_*LFI*_ is the locally optimal linear estimator [27].

The Shannon mutual information quantifies the reduction in uncertainty of one random variable given knowledge of another

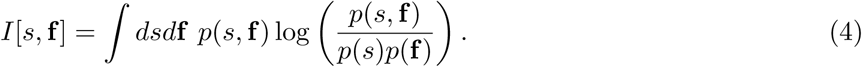

Earlier work demonstrated that the Fisher information provides a lower bound for the Shannon mutual information in the case of Gaussian noise [11]. However, more recent work has revealed that the relationship between the two is more nuanced, particularly in the cases where the noise model is non-Gaussian [47]. Thus, we supplement our assessment of the network’s coding ability by measuring the mutual information, *I*[*s*, **r**], between the neural representation **r** and the stimulus *s*. As with the Fisher information, the mutual information is often intractable, but fortunately can be estimated from data. Specifically, we will employ the estimator developed by Kraskov and colleagues, which utilizes entropy estimates from *k*-nearest neighbor distances [28].

### 2.3 Structured Weights

The measures of coding strength are a function of the weights that shape the interaction of the stimulus and noise in the network. Thus, the choice of the synaptic weight distribution impacts the calculation of these quantities. We first consider the case of “structured weights” in order to obtain analytical expressions for measures of coding strength. Structured weights take on the form

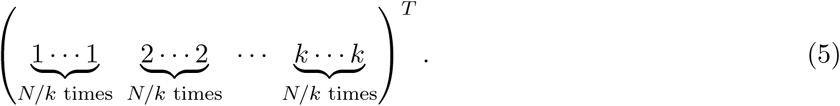

Specifically, the structured weight vectors are parameterized by an integer *k* which divides the *N* weights into *k* homogeneous groups. The weights across the groups span the positive integers up to *k*. Importantly, larger *k* will only increase the weights in the vector. Thus, in the above scheme, increased “diversity” can only be achieved by increasing *k*, which will invariably result in an amplification of the signal to which the weight vector is applied. In the case that *k* does not evenly divide *N*, each group is repeated ⌈*N/k*⌉ times, except the last group, which is only repeated *N −* (*N −* 1) ⋅ ⌈*N/k*⌉ times (*i.e.*, the last group is truncated to ensure the weight vector is of size *N*).

Additionally, we consider cases in which *k* is of order *N*, *e.g.*, *k* = *N/*2. Allowing *k* to grow with *N* ensures that typical values for the weights grow with the population size. This contrasts with the case in which *k* is a constant, such as *k* = 4, which sets a maximum weight value independent of the population size.

### 2.4 Unstructured Weights

While the structured weights allow for analytical results, they possess an unrealistic distribution of synaptic weighting. Thus, we also consider the case of “unstructured weights,” in which the synaptic weights are drawn from some parameterized probability distribution:

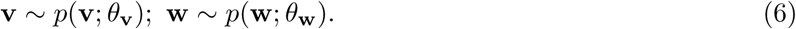

We calculate both information theoretic quantities over many random draws from these distributions, and observe how these quantities behave as some subset of the parameters *θ* are varied. In particular, we focus on the lognormal distribution, which has been found to describe the distribution of synaptic weights well in slice electrophysiology [40, 45]. Specifically, the weights take on the form

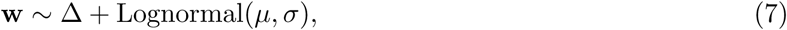

where ∆ *>* 0. For a lognormal distribution, an increase in *µ* will increase the distribution’s mean, median, and mode (Fig. 3e, inset). Thus, *µ* as a parameter acts similarly to *k* for the structured weights in that increased weight diversity must be accompanied by an increase in their magnitude.

## 3 Results

We consider the network’s coding ability after both the linear stage (ℓ) and the nonlinear stage (**r**). In other words, the linear stage can be considered the output of the network assuming each of the functions *g*_*i*_(ℓ_*i*_) is the identity. Furthermore, due to the data processing inequality, the qualitative conclusions we obtain from the linear stage should apply for any one-to-one nonlinearity.

**Figure 3:**
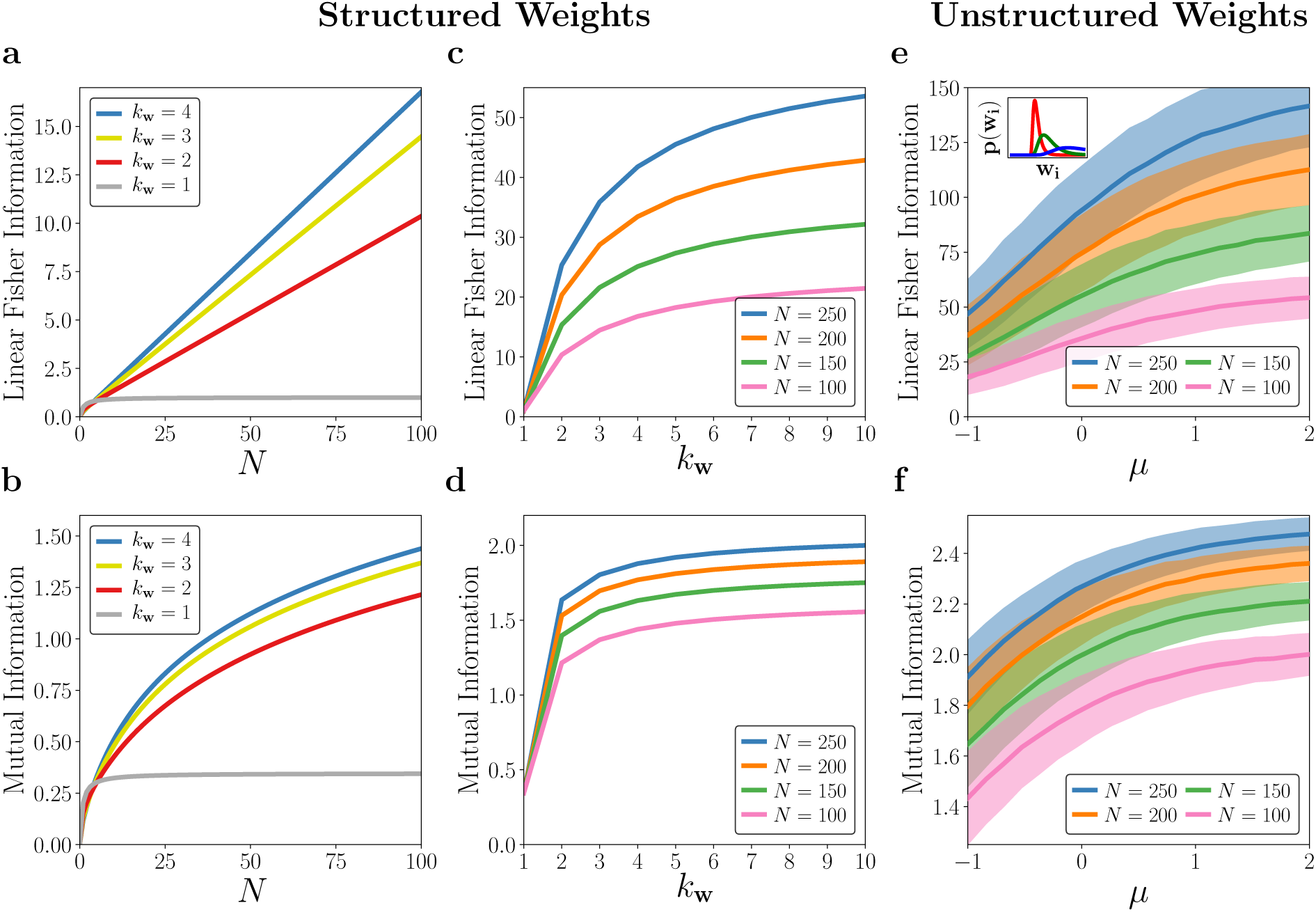
Network coding performance of the linear stage representation. Here, the noise variances are 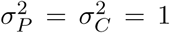. Fisher information is shown on the top row while mutual information is shown on the bottom row. **(a)**, **(b)** Structured weights. Linear Fisher Information and Mutual Information are shown as a function of the population size, *N*, across different levels of weight heterogeneity, *k*_**w**_ (indicated by color). **(c)**, **(d)** Linear Fisher Information and Mutual Information are shown as a function of weight heterogeneity, *k*_**w**_, for various population sizes, *N*. **(e)**, **(f)** Unstructured weights. Linear Fisher Information and Mutual Information are shown as a function of the mean of the lognormal distribution used to draw common noise synaptic weights. Information quantities are calculated across 1000 random drawings of weights: solid lines depict the means while the shaded region indicates one standard deviation. Inset: the distribution of weights for various choices of *µ*. Increasing *µ* shifts the distribution to the right, increasing heterogeneity.

### 3.1 Linear Stage

The Fisher information about the stimulus in the linear representation can be shown to be (see Appendix 5.1 for the derivation)

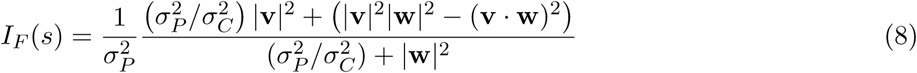

which is equivalent to the linear Fisher information in this case. The mutual information can be expressed as (see Appendix 5.2 for the derivation)

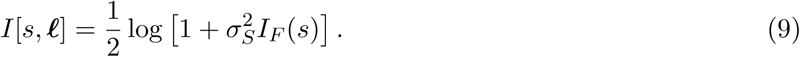

For the case the mutual information, we have assumed the prior distribution for the stimulus is Gaussian with zero mean and variance 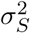. For structured weights, equations (8) and (9) can be explored by varying the choice of *k* for both **v** and **w** (we will refer to them as *k*_**v**_ and *k*_**w**_, respectively).

It is simplest and most informative to examine these quantities by setting *k*_**v**_ = 1 while allowing *k*_**w**_ to vary, as amplifying and diversifying **v** will only increase coding ability for predictable reasons (this is indeed the case for our network) [17, 42]. While increasing *k*_**w**_ will boost the overall amount of noise added to the neural population, it also changes the direction of the noise in the higher-dimensional neural space. Thus, while we might expect that adding more noise in the system would hinder coding, the relationship between the directions of the noise and stimulus vectors in the neural space also plays a role.

We first consider how the Fisher information and mutual information are impacted by the choice of *k*_**w**_. In the structured regime, we have

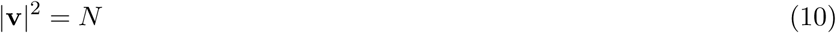

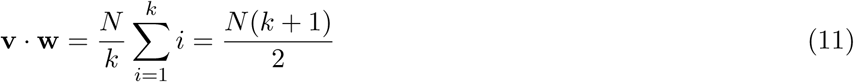

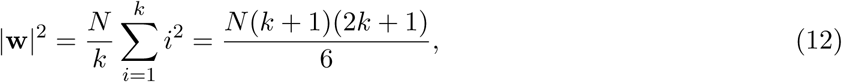

which allows us to rewrite equation (8) as

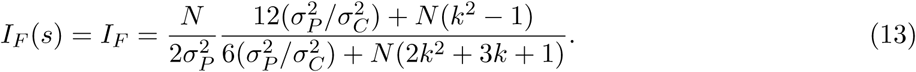

The form of the mutual information follows directly from plugging equation (13) into equation (9).

The analytical expressions for the structured regime reveal the asymptotic behavior of the information quantities. Neither quantity saturates as a function of the number of neurons, *N*, except in the case of *k*_**w**_ = 1 (Fig. 3a, b). In this regime, increasing the population size of the system also enhances coding fidelity. Furthermore, both quantities are monotonically increasing functions of the common noise synaptic heterogeneity, *k*_**w**_ (Fig. 3c, d), implying that decoding is enhanced despite the fact that the amplitude of the common noise is magnified for larger *k*_**w**_. Our analytical results show linear and logarithmic growth for the Fisher and mutual information, respectively, as one might expect in the case of Gaussian noise [11]. These qualitative results hold for essentially any choice of (*σ*_*S*_, *σ*_*P*_, *σ*_*C*_).

In the case of *k*_**w**_ = 1, the signal and common noise are aligned perfectly in the neural representation. Thus, the common noise becomes equivalent in form to shared input noise. As a consequence, we observe the saturation of both Fisher information and mutual information as a function of the neural population. This saturation implies the existence of differential correlations, consistent with the observation that information-limiting correlations occur under the presence of shared input noise [24].

The structured weight distribution described above allows us to derive analytical results, but the limitation to only a fixed number of discrete synaptic weight values is not realistic for biological networks. Thus, we utilize unstructured weights, described in Section 2.4, in which the synaptic weights are drawn from a lognormal distribution. In this case, we estimate the linear Fisher information and the mutual information over many random draws according to *w*_*i*_ *∼* ∆+Lognormal(*µ, σ*^2^). We are primarily concerned with varying *µ*, as an increase in this quantity uniformly increases the mean, median, and mode of the lognormal distribution (Fig. 3e, inset), akin to increasing *k*_**w**_ for the structured weights.

Our numerical analysis demonstrates that increasing *µ* increases the average Fisher information and average mutual information across population sizes (Fig. 3e, f: bold lines). In addition, the benefits of larger weight diversity are felt more strongly by larger populations (Fig. 3e, f: different colors).

In the structured weight regime, our analytical results show that weight heterogeneity can ameliorate the harmful effects of *additional* information-limiting correlations induced by common noise mimicking shared input noise. They do not imply that weight heterogeneity prevents differential correlations, as the common noise in this model is manipulated by synaptic weighting, in contrast with true shared input noise. For unstructured weights, we once again observe that larger heterogeneity affords the network improved coding performance, despite the increased noise in the system. Together, these results show that linear networks can manipulate common noise to prevent it from causing differential correlations.

### 3.2 Quadratic Nonlinearity

We next consider the performance of the network after a quadratic nonlinearity *g*_*i*_(*x*) = *x*^2^ for all neurons *i* [35]. In this case, both the Fisher information and mutual information are analytically intractable. Thus, we will instead turn to the linear Fisher information, which can be calculated, and approximate the mutual information numerically.

#### 3.2.1 Linear Fisher Information

An analytic expression of the linear Fisher information is calculated in Appendix 5.3. Its analytic form is too complicated to be restated here, but we will examine it numerically for both the structured and unstructured weights. The qualitative behavior of the Fisher information depends on the magnitude of the common variability (*σ*_*C*_) and private variability (*σ*_*P*_) in a more complicated fashion than the linear stage, which depends on these variables primarily through their ratio *σ*_*C*_/*σ*_*P*_. Thus, we separately consider how common and private variability impact coding efficacy under various synaptic weight structures.

As before, we first consider the structured weights with *k*_**v**_ set to 1 while only varying *k*_**w**_. We start with the special case where *σ*_*P*_ = *σ*_*C*_ = 1 (*i.e.*, equal private and common noise variance). Here, the Fisher information saturates for both *k*_**w**_ = 1 and *k*_**w**_ = 2, but increases without bound for larger *k*_**w**_ (Fig. 4a). We can also consider the case where the structured weight heterogeneity grows in magnitude with the population size (*i.e.*, *k*_**w**_ is a function of *N*). In this scenario, the Fisher information is much smaller and saturates (Fig. 4a, dashed lines). This implies the existence of differential correlations.

**Figure 4:**
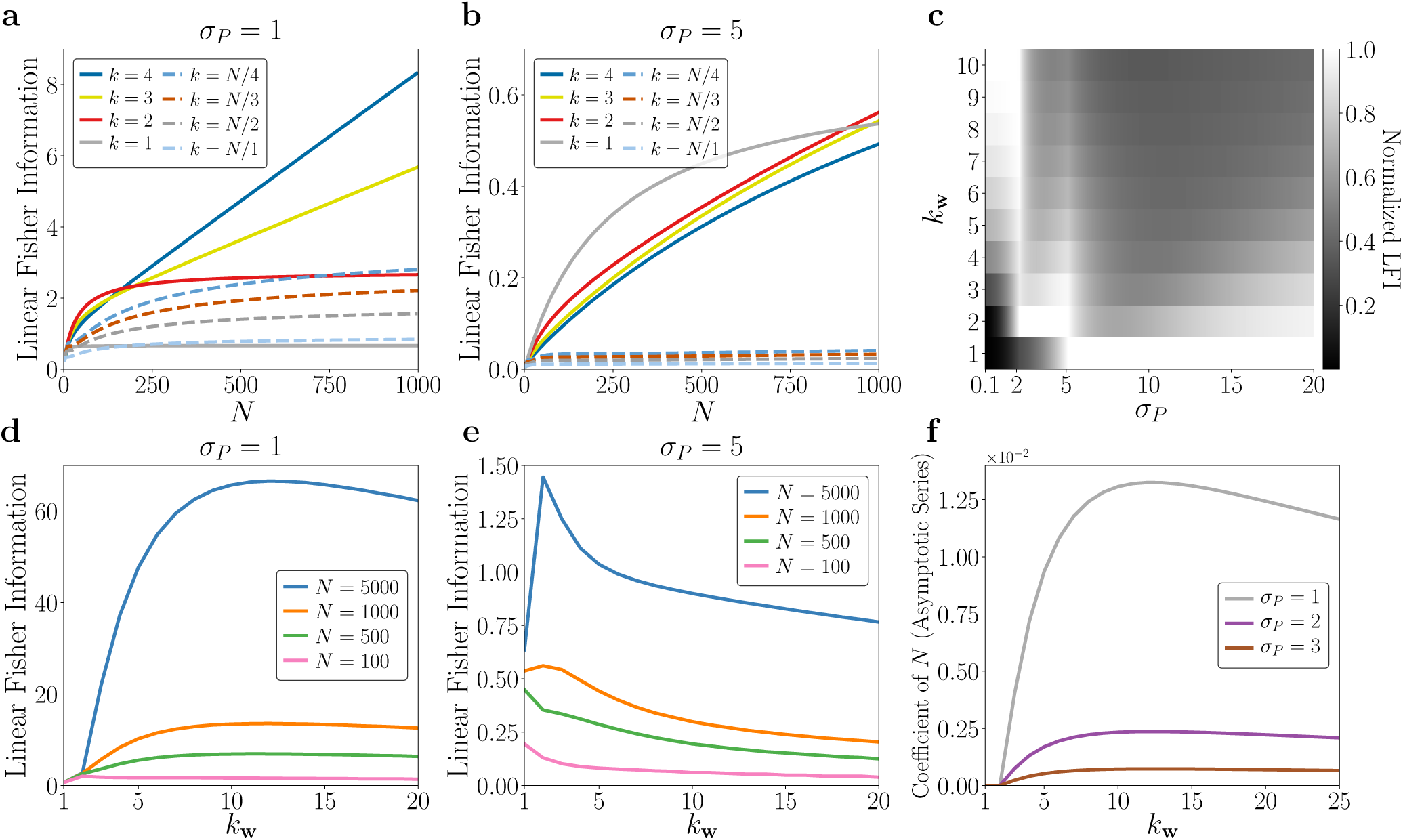
Linear Fisher information after quadratic nonlinearity in a network with structured weights. **(a)** Fisher information as a function of population size when *σ*_*P*_ = *σ*_*C*_ = 1, *i.e.*, private and common noise have equal variances. Solid lines denote constant *k* while dashed lines denote *k* scaling with population size. **(b)** Same as (a), but for a network where private variance dominates (*σ*_*P*_ = 5, *σ*_*C*_ = 1). **(c)** Normalized fisher information: for a choice of *σ*_*P*_, the Fisher information is calculated for a variety of *k*_**w**_ (*y*-axis) and divided by the maximum Fisher information (across the *k*_**w**_, for the choice of *σ*_*P*_). For a given *σ*_*P*_, the normalized Fisher information is equal to one at the value of *k*_**w**_ which maximizes decoding performance. **(d)** Behavior of the Fisher information as a function of synaptic weight heterogeneity for various population sizes (*σ*_*P*_ = *σ*_*C*_ = 1). **(e)** Same as (d), but for networks where private variance dominates (*σ*_*P*_ = 5, *σ*_*C*_ = 1). **(f)** The coefficient of the linear term in the asymptotic series of the Fisher information at different levels of private variability. At *k*_**w**_ = 1, 2, the coefficient of *N* is exactly zero.

When private variability dominates, we observe qualitatively different finite network behavior (*σ*_*P*_ = 5, Fig. 4b). For *N* = 1000, both *k*_**w**_ = 1 and *k*_**w**_ = 2 exhibit better performance relative to larger values of *k*_**w**_ (by contrast, the case with *k*_**w**_ *∼ O*(*N*) quickly saturates). We note that, unsurprisingly, the increase in private variability has decreased the Fisher information for all cases we considered compared to *σ*_*P*_ = 1 (compare the scales of Fig. 4a and Fig. 4b). Our main interest, however, is identifying effective synaptic weighting strategies *given* some amount of private and common variability.

The introduction of the squared nonlinearity produces qualitatively different behavior at the finite network level: in contrast with Figure 3, increased heterogeneity does not automatically imply improved decoding. In fact, there is a regime in which increased heterogeneity improves Fisher information, beyond which we see a reduction in decoding performance (Fig. 4d). If the private variability is increased, this regime shrinks or becomes nonexistent, depending on the population size (Fig. 4e). Furthermore, entering this regime for higher private variability requires smaller *k*_**w**_ (*i.e.*, less weight heterogeneity).

The results shown in Figure 4d and Figure 4e imply that there exists an interesting relationship between the network’s decoding ability, its private variability, and its synaptic weight heterogeneity *k*_**w**_. To explore this further, we examine the behavior of the Fisher information at a fixed population size (*N* = 1000) as a function of both *σ*_*P*_ and *k*_**w**_ (Fig. 4c). To account for the fact that an increase in private variability will always decrease the Fisher information, we calculate the *normalized* Fisher information: for a given choice of *σ*_*P*_, each Fisher information is divided by the maximum across a range of *k*_**w**_ values. Thus, a normalized Fisher information allows us to determine what level of synaptic weight heterogeneity maximizes coding fidelity, given a particular level of private variability *σ*_*P*_.

Figure 4c highlights three interesting regimes. When the private variability is small, the network benefits from larger weight heterogeneity on the common noise. But as the neurons become more noisy, the “Goldilocks zone” in which the network can leverage larger noise weights becomes constrained. When the private variability is large, the network is achieves superior coding fidelity by having less heterogeneous weights, despite the threat of induced differential correlations from the common noise. Between these regimes, there are transitions for which many choices of *k*_**w**_ result in equally good decoding performance.

It is important to point out that Figures 4a-e only captures finite network behavior. Therefore, we extended our analysis by validating the asymptotic behavior of the Fisher information as a function of the private noise by examining its asymptotic series at infinity (Fig. 4f). For *k*_**v**_ = 1, 2, the coefficient of the linear term is zero for any choice of *σ*_*P*_, implying that the Fisher information always saturates. In addition, when the common noise weights increase with population size (*i.e.*, *k*_**w**_ *∼ O*(*N*)), the asymptotic series is always sublinear (not shown in Fig. 4f). Thus, there are multiple cases in which the structure of synaptic weighting can induce differential correlations in the presence of common noise. Increased heterogeneity allows the network to escape these induced differential correlations and achieve linear asymptotic growth. If *k*_**w**_ becomes too large, however, the linear asymptotic growth begins to decrease. Once *k*_**w**_ scales as the population size, differential correlations are once again significant.

Next, we reproduce the above analysis with unstructured weights. As before, we draw 1000 samples of common noise weights from a shifted lognormal distribution with varying *µ*. The behavior of the average (linear) Fisher information is qualitatively similar to that of the structured weights (Fig. 5). There exists a regime for which larger weight heterogeneity improves the decoding performance, beyond which coding fidelity decreases (Fig. 5a). If the private noise variance dominates, this regime begins to disappear for smaller networks (Fig. 5b). Thus, with very noisy neurons, the coding fidelity of the network is improved when the synaptic weights are less heterogeneous (and therefore, smaller).

**Figure 5:**
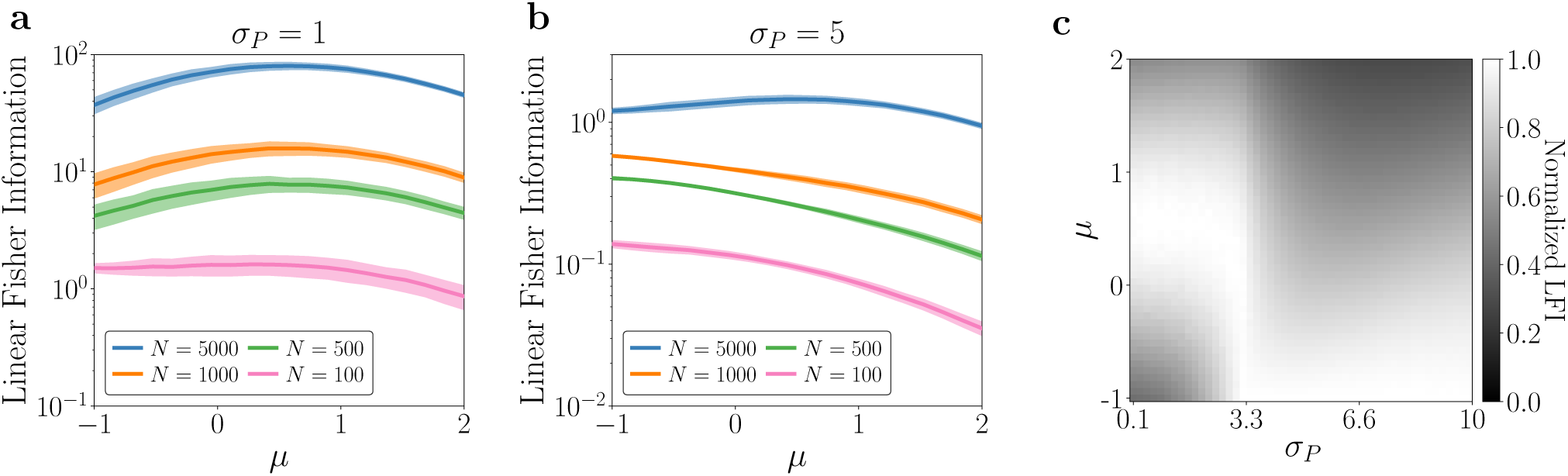
Linear Fisher information after quadratic nonlinearity, unstructured weights. In contrast to Figure 4, subplots (a) and (b) are plotted on a log-scale. **(a)** Linear Fisher information as a function of the mean, *µ*, of the lognormal distribution used to draw the common noise synaptic weights. Solid lines denote means while shaded regions denote one standard deviation across the 1000 drawings of weights from the lognormal distribution. **(b)** Same as (a), but for networks in which private variability dominates (*σ*_*P*_ = 5, *σ*_*C*_ = 1). **(c)** Normalized Linear Fisher information. Same plot as Figure 4c, but the average Fisher information across the 1000 samples is normalized across *µ* (akin to normalizing across *k*_**w**_).

To summarize these results, we once again plot the normalized Fisher information (this time, normalized across choices of *µ* and averaged over 1000 samples from the lognormal distribution) for a range of private variabilities (Fig. 5c). The heat map exhibits a similar transition at a specific level of private variability. At this transition, a wide range of *µ*’s provide the network with similar decoding ability. For smaller *σ*_*P*_, we see behavior comparable to Figure 5a, where there exists a regime of improved Fisher information. Beyond the transition, the network performs better with less diverse synaptic weighting, though it becomes less stringent as *σ*_*P*_ increases. The behavior exhibited by this heat map is similar to Figure 4c, but contains fewer uniquely identifiable regions. This may imply that the additional regions in Figure 4c are an artifact of the structured weights.

The amount of the common noise will also impact how the network behaves and what levels of synaptic weight heterogeneity are optimal. For example, consider a network with private noise variability set to *σ*_*P*_ = 1. When common noise is small, the Fisher information is comparable among various choices of synaptic weight diversity (Fig. 6a). When the common noise dominates, however, the network benefits strongly from diverse weighting (Fig. 4b), though it is punished less severely for having *k*_**w**_ scale with *N* (Fig. 6b, dashed lines; compare to Fig. 4b). These observations are true at finite population size. As before, the Fisher information saturates for *k*_**w**_ = 1, 2 and *k*_**w**_ *∼ O*(*N*), no matter the choice of common noise variance.

**Figure 6:**
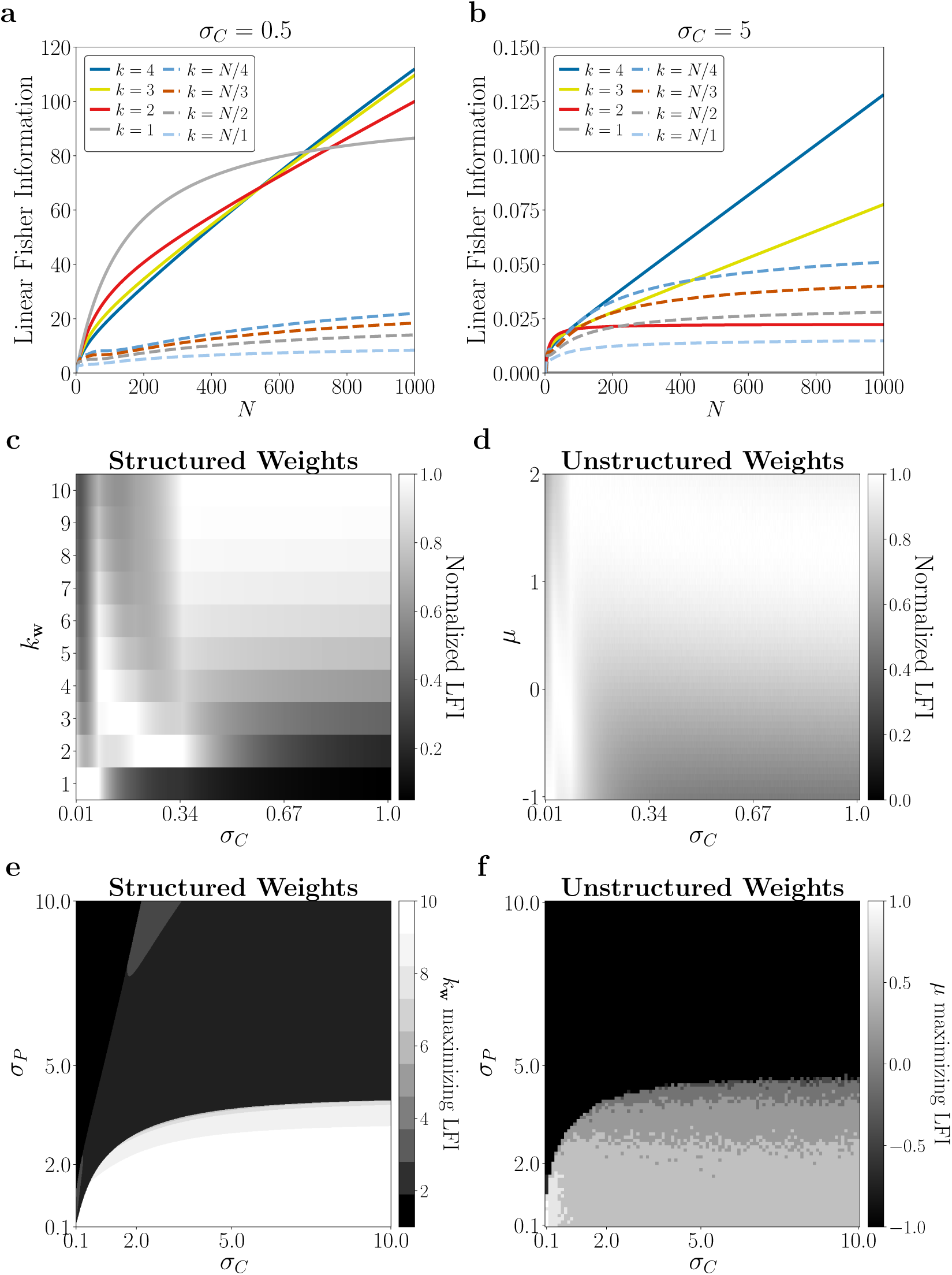
The relationship between common noise, private noise, and synaptic weight heterogeneity. **(a), (b)** Fisher information as a function of population size, *N*, when common noise contribution is drowned out by private noise (a), and when common noise dominates (*σ*_*P*_ = 1) (b). Solid lines indicate constant *k*_**w**_ while dashed lines refer to *k*_**w**_ that scales with *N*. **(c), (d)** Normalized Fisher information as a function of common noise for structured weights (c) and unstructured weights (d). For unstructured weights, each Fisher information is calculated by averaging over 1000 networks with their common noise weights drawn from the respective distribution. **(e)** The value of *k*_**w**_ that maximizes the network’s Fisher information for a given choice of *σ*_*P*_ and *σ*_*C*_. The maximum is taken over over *k*_**w**_ *∈* [1, 10]. **(f)** The value of *µ* that maximizes the average Fisher information over 1000 draws for a given choice of *σ*_*P*_ and *σ*_*C*_.

We calculated the normalized Fisher information across a range of common noise strengths to determine the optimal synaptic weight distribution. The results for structured weights and unstructured weights are shown in Figures 6c and 6d, respectively. While they strongly resemble Figure 4c and Figure 5c, they exhibit opposite qualitative behavior. As before, there are three identifiable regions in Figure 6c, each divided by abrupt transitions where many choices of *k*_**w**_ are equally good for decoding. For small common noise, the coding fidelity is improved with less heterogeneous weights, but as the common noise increases, the network enters the “Goldilocks regions”. After another abrupt transition near *σ*_*C*_ *≈* 0.34, the network performance is greatly improved by heterogeneous weights.

Thus, common noise and private noise seem to have opposite impacts on the optimal choice of synaptic weight heterogeneity. When private noise dominates, the Fisher information is maximized under a set of homogeneous weights, since coding ability is harmed by amplification of common noise. When common noise dominates, the network coding is improved under diverse weighting: this prevents additional differential correlations and furthermore helps the network cope with the punishing effects on coding due to the amplified noise.

How should we choose the synaptic weight distribution within the extremes of private or common noise dominating? We assess the behavior of the Fisher information as both *σ*_*P*_ and *σ*_*C*_ are varied over a wide range. For the structured weights, we calculate the choice of *k*_**w**_ that maximized the network’s Fisher information (within the range *k*_**w**_ *∈* [1, 10]) (Fig. 6e). For the unstructured weights, we calculated the choice of *µ* that maximizes the network’s average Fisher information over 1000 drawings of **w** from the lognormal distribution specified by *µ* (Fig. 6f).

Figures 6e and 6f reveal that the network is highly sensitive to the values of *σ*_*P*_ and *σ*_*C*_. Figure 6e exhibits a band like structure and abrupt transitions in the value of *k*_**w**_ which maximizes Fisher information. This band-like structure would most likely continue to form for smaller *σ*_*P*_ if we allowed *k*_**w**_ *>* 10. One might expect that the band-like structure is due to the artificial structure in the weights; however, we see that Figure 6f also exhibits these types of bands. Note that the regime of interest for us is when private variability is a smaller contribution to the total variability than the common variability. When this is the case, Figures 6e and 6f imply that a population of neurons will be best served by having a diverse set of synaptic weights, even if the weights amplify irrelevant signals.

Together, these results highlight how the introduction of the nonlinearity in the network reveal an intricate relationship between the amount of shared variability, private variability, and the optimal synaptic weight heterogeneity. Our observations that the network benefits from increased synaptic weight heterogeneity in the presence of common noise are predicated on the size of the network (Fig. 4a-b, Fig. 6a-b) and the amount of private and shared variability (Fig. 4c, Fig 6c-d). In particular, when shared variability is the more significant contribution to the overall variability, the coding performance of the network benefits from increased heterogeneity, whether the weights are structured or unstructured (Fig. 6e-f). This implies that, in contrast to the linear network, there exist regimes where increasing the synaptic weight heterogeneity beyond a point will harm coding ability (Fig. 4d-e, Fig 5a-b), demonstrating that there is a tradeoff between the benefits of synaptic weight heterogeneity and the amplification of common noise it may introduce.

#### 3.2.2 Mutual Information

When the network possesses a quadratic nonlinearity, the mutual information *I*[*s*, **r**] is far less tractable than for the linear case. Therefore, we computed the mutual information numerically on data simulated from the network, using an estimator built on *k*-nearest neighbor statistics [28]. We refer to this estimator as the KSG estimator.

We applied the KSG estimator to 100 unique datasets, each containing 100,000 samples drawn from the linear-nonlinear network. We then estimated the mutual information within each of the 100 datasets. The computational bottleneck for the KSG estimator lies in finding nearest neighbors in a *kd*-tree, which becomes prohibitive for large dimensions (*∼*20), so we considered much smaller population sizes than in the case of Fisher information. Furthermore, the KSG estimator encountered difficulties when samples became too noisy, so we limited our analysis to smaller values of (*σ*_*P*_, *σ*_*C*_). Due to these constraints, we are only able to probe the finite network behavior of the mutual information.

Our results for the structured weights are shown in Figure 7. When utilizing estimators of mutual information from data, caution should be taken before comparing across different dimensions, due to bias in the KSG estimator [20]. Thus, we restrict our observations to within a specified population size. First, we evaluated the mutual information for various population sizes (*N* = 8, 10, 12, 14) in the case where *σ*_*C*_ = *σ*_*P*_ = 0.5. Observe that, as before, the mutual information increases with larger weight heterogeneity (*k*_**w**_, Fig. 7a). The improvement in information occurs for all four population sizes.

**Figure 7:**
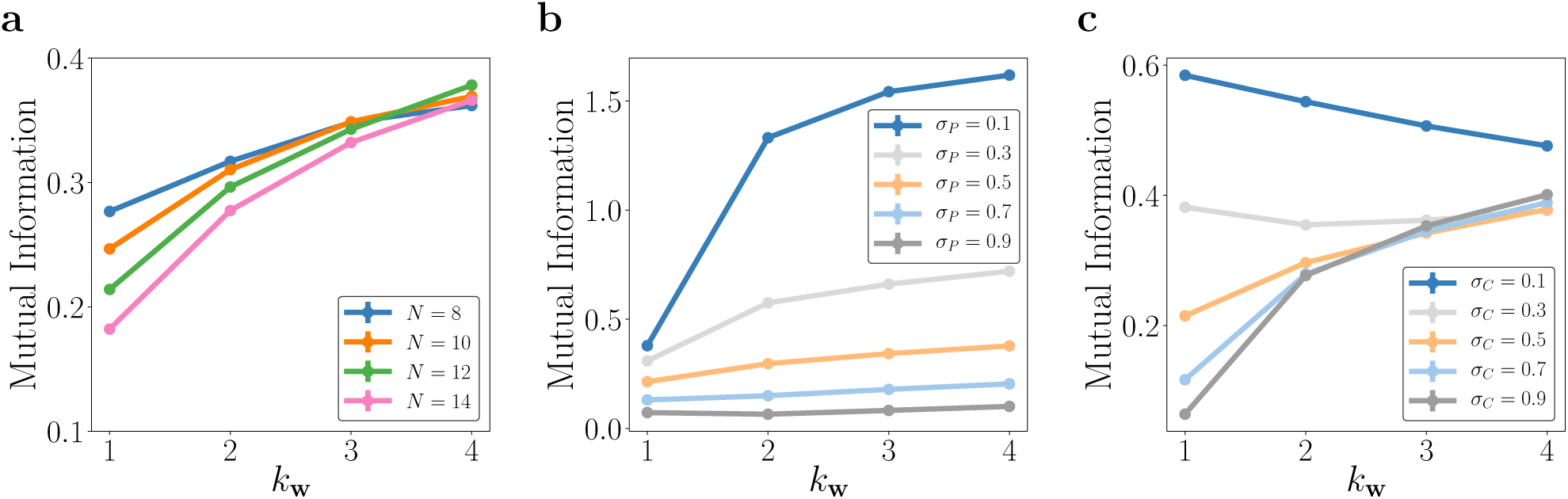
Mutual information computed by applying the KSG estimator on data simulated from the network with quadratic nonlinearity and structured weights. The estimates consist of averages over 100 datasets, each containing 100,000 samples. Standard error bars are smaller than the size of the markers. **(a)** Mutual information as a function of common noise weight heterogeneity for various population sizes *N*. We consider smaller *N* than in the case of Fisher information as computation time becomes prohibitive for larger dimensionalities. Here, *σ*_*P*_ = *σ*_*C*_ = 0.5. **(b)** The behavior of mutual information for various choices of *σ*_*P*_, while *σ*_*C*_ = 0.5. **(c)** The behavior of mutual information for various choices of *σ*_*C*_, while *σ*_*P*_ = 0.5.

Decreasing the private variability increases mutual information (Fig. 7b). However, the network sees a greater increase in information with diverse weighting when *σ*_*P*_ is small. This is consistent with the small *σ*_*P*_ regime highlighted in Figure 4c: the smaller the private variability, the more the network benefits from larger synaptic weight heterogeneity. Similarly, decreasing the common variability increases mutual information (Fig. 7c). If the common variability is small enough (for example, *σ*_*C*_ = 1), then larger *k*_**w**_ harms the encoding. Thus, when the common noise is small enough, the amplification of noise that results when *k*_**w**_ is increased harms the network’s encoding. It is only when the common variability becomes the dominant contribution to the variability that the diversification provided by larger *k*_**w**_ improves the mutual information.

As for the unstructured weights, we calculated the mutual information *I*[*s*, **r**] over 100 synaptic weight distributions drawn from the aforementioned lognormal distribution. For each synaptic weight distribution, we applied the KSG estimator to 100 unique datasets, each consisting of 10,000 samples. Thus, the mutual information estimate for a given network was computed by averaging over the individual estimates across the 100 datasets. With this procedure, we explored how the mutual information behaves as a function of the private noise variability, common noise variability, and mean of the lognormal distribution.

Similar to the normalized Fisher information, we present the normalized mutual information as a function of the private and common variances (Fig. 8). For a given *σ*_*P*_ or *σ*_*C*_, the mutual information is calculated across a range of *µ ∈* [*−*1, 1]. The normalized mutual information is obtained by dividing each individual mutual information by the maximum value across the *µ*. Thus, for a given *σ*_*P*_, the value of *µ* whose normalized mutual information is 1 specifies the lognormal distribution that maximizes the network’s encoding performance. As private variability increases, the network benefits more greatly benefits diverse weighting (larger *µ*, Fig. 8a). As common variability increases, the network once again prefers more diverse weighting. If the common variability is small enough, however, the network is better suited to homogeneous weights (Fig. 8b). Therefore, the analysis utilizing the unstructured weights largely corroborates our findings for the structured weights shown in Figure 7.

**Figure 8:**
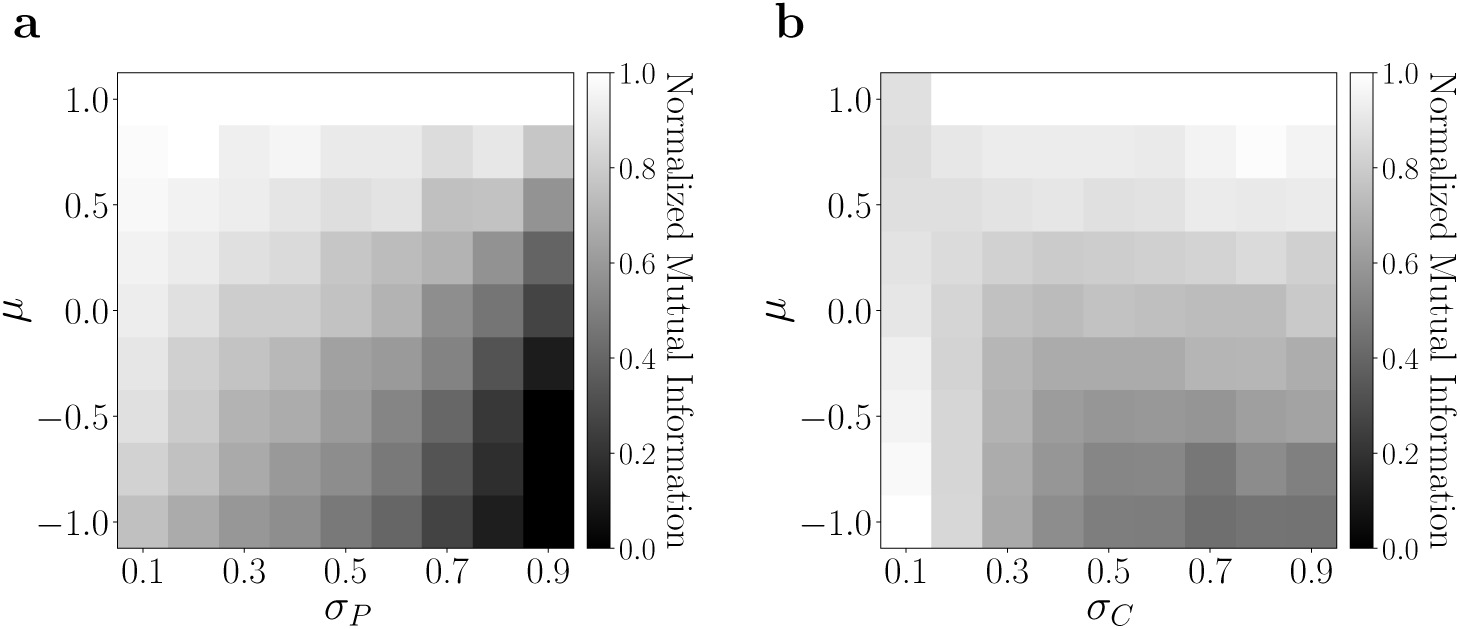
Normalized mutual information for common and private variability. For a given *µ*, 100 networks were created by drawing common noise weights **w** from the corresponding lognormal distribution. The mutual information shown is the average across the 100 networks. For a specified network, the mutual information was calculated by averaging KSG estimates over 100 simulated datasets, each containing 10,000 samples. Finally, for a choice of (*σ*_*P*_, *σ*_*C*_), mutual information is normalized to the maximum across values of *µ*. **(a)** Normalized mutual information as a function of *µ* and private variability (*σ*_*C*_ = 0.5). **(b)** Normalized mutual information as a function of *µ* and common variability (*σ*_*P*_ = 0.5).

Thus, these results highlight that there exist regimes where neural coding, as measured by the Shannon mutual information, benefit from increased synaptic weight heterogeneity. Furthermore, similarly to the case of the linear Fisher information, the improvement in coding occurs more significantly when shared variability is large relative to private variability.

## 4 Discussion

We have demonstrated in a simple model of neural activity that if synaptic weighting of common noise inputs is broad and heterogeneous, coding fidelity is actually improved despite inadvertent amplification of common noise inputs. We showed that for squaring nonlinearities, there exists a regime of heterogeneous weights for which coding fidelity is maximized. We also found that the relationship between the magnitude of private and shared variability is vital for determining the ideal amount of synaptic heterogeneity. In neural circuits where shared variability is dominant, as has been reported in some parts of the cortex [16], larger weight heterogeneity results in better coding performance (Fig. 6e).

Why are we afforded improved neural coding under increased synaptic weight heterogeneity? An increase in heterogeneity, as we have defined it, ensures that the common noise is magnified in the network. At the same time, however, the structure of the correlated variability induced by the common noise is altered by increased heterogeneity. Previous work has demonstrated that the relationship between signal correlations and noise correlations is important in assessing decoding ability: for example, the sign rule states that noise correlations are beneficial if they are of opposite sign as the signal correlation [22]. Geometrically, the sign rule is a consequence of the intuitive observation that decoding is easier when the noise correlations lie perpendicular to the signal manifold [4, 33, 53].

For example, consider the correlated activity for two neurons in the network against their signal space (black lines, Fig. 9a, b) as a function of *k*_**w**_. Note that the signal space is linear, due to the quadratic linearity (see Appendix). After the linear stage, the larger weight heterogeneity pushes the cloud of neural activity to lie more orthogonal to the signal space. At the same time, the variance becomes observably larger due to the magnification of the common noise (Fig. 9a). Importantly, note that the variability for *k*_**w**_ = 1 lies parallel to the signal space, signifying the presence of differential correlations. The correlated variability after the nonlinear stage is similar in that orthogonality to the signal space increases with *k*_**w**_. There is a notable difference: squaring the linear stage ensures non-negative activities, thereby limiting the response space. Thus, for large enough *k*_**w**_, the rectification manifests strongly enough that the network enters a regime where increased heterogeneity harms decoding. These figures only demonstrate the relationship between a pair of neurons, while the collective correlated variability structure ultimately dictates decoding performance. They do, however, shed light on how the distribution of synaptic weights can radically shape the common noise and thereby the overall structure of the shared variability.

**Figure 9:**
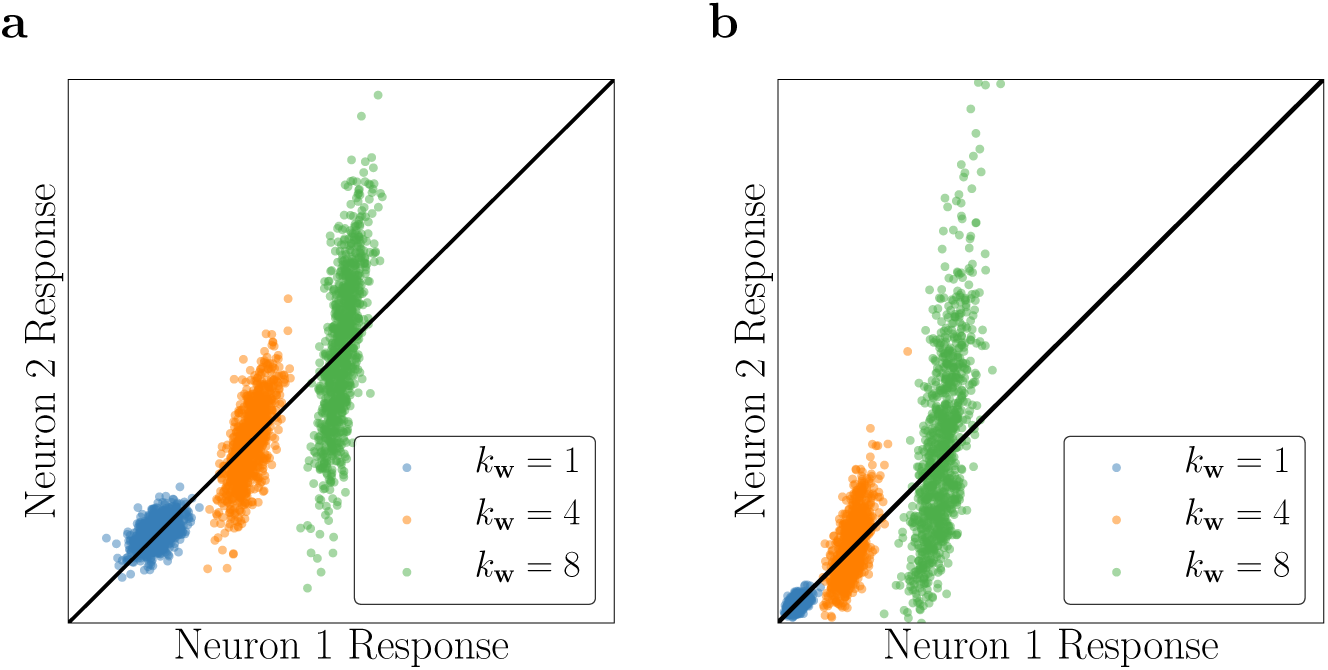
The benefits of increased synaptic weight heterogeneity. **(a)** The responses of a pair of neurons against the signal space, taken after the linear stage. Colors indicate different choices of *k*_**w**_ (while *k*_**v**_ = 1). Each cloud contains 1000 sampled points. **(b)** Same as (a), but responses are taken after the quadratic nonlinearity.

The linear stage of the network constitutes a noisy projection of two signals (one of which is not useful to the network) in a high-dimensional space. Thus, we can assess the entire population by examining the relationship between the projecting vectors **v** and **w**. We might expect that improved decoding occurs when these signals are farther apart in the *N*-dimensional space [23]. For a chosen *k*_**v**_, this occurs as *k*_**w**_ is increased when the weights are structured. When the weights are unstructured, the average angle between the stimulus and weight vectors is large as either *µ*_*v*_ or *µ*_*w*_ increases. Increased heterogeneity implies access to a more diverse selection of weights, thus pushing the two signals apart. From this perspective, the nonlinear stage acts as a mapping on the high-dimensional representation. Given that no noise is added after the nonlinear processing stage in the networks, if the nonlinearities were one-to-one, the data processing inequality would ensure that the results from the linear stage would hold. But, as we observed earlier, the nonlinear stage benefits from increased heterogeneity only in certain regimes. Thus, the rectifying nature of the nonlinearities is important: the application of both the quadratic nonlinearity restricts the high-dimensional space that the neural code can occupy, and thus limits the benefits of diverse synaptic weighting. We would expect similar behavior if the neural activity were passed through a Poisson distribution, further rectifying the outputs.

It may seem unreasonable that the neural circuit possesses the ability to weight common noise inputs. However, excitatory neurons receive many excitatory synapses in circuits throughout the brain. Some subset of common inputs across a neural population will undoubtedly be irrelevant for the underlying neural computation, even if these signals are not strictly speaking “noise” and could be useful for other computations. Thus, these populations must contend with common noise sources contributing to their overall shared variability and potentially hampering their ability to encode a stimulus. Our work demonstrates that neural circuits, armed with a good set of synaptic weights, need not suffer adverse impacts due to inadvertently amplifying potential sources of common noise. Instead, broad, heterogeneous weighting ensures that common noise sources will project the signal and noise into a high-dimensional space in such a way that is beneficial for decoding. This observation is in agreement with recent work that explored the relationship between heterogeneous weighting and degrees of synaptic connectivity [31]. Furthermore, synaptic input, irrelevant on one trial, may become the signal on the next: heterogeneous weighting provides a general, robust principle for neural circuits to follow.

We chose the simple network architecture in order to maintain analytic tractability, which allowed us to explore the rich patterns of behavior it exhibited. Our model is limited, however. It is worthwhile to assess how our qualitative conclusions hold with added complexity in the network. For example, interesting avenues to consider include the implementation of recurrence, spiking dynamics, and other, thresholded nonlinearities (*e.g.*, rectified linear unit or a squared threshold). In addition, these networks could also be equipped with varying degrees of sparsity and inhibitory connections. Importantly, the balance of excitation and inhibition in networks has been shown to be vital in decorrelating neural activity [38]. Past work has explored how to approximate both information theoretic quantities studied here in networks with some subset of these features [8, 50]. Thus, analyzing how common noise and synaptic weighting interact in more complex networks is of interest for future work.

We established correlated variability structure in the linear-nonlinear network by taking a linear combination of a common noise source and private noise sources (though our model ignores any noise potentially carried by the stimulus). This was sufficient to establish low-dimensional shared variability observed in neural circuits. As a consequence, our model as devised enforces stimulus-independent correlated variability. Recent work, however, has demonstrated that correlated variability is in fact stimulus-dependent. Such work used both phenomenological [19, 30] and mechanistic [53] models in producing fits to the stimulus-dependent correlated variability. These models all share a doubly stochastic noise structure, stemming from both additive and multiplicative sources of noise [21]. It is therefore worthwhile to fully examine how both additive and multiplicative sources of noise interact with synaptic weighting to influence neural coding.

## Acknowledgments

We thank Ruben Coen-Cagli for useful discussions. P. S. S. was supported by the Department of Defense (DoD) through the National Defense Science & Engineering Graduate Fellowship (NDSEG) Program. J. A. L. was supported through the Lawrence Berkeley National Laboratory-internal LDRD “Deep Learning for Science” led by Prabhat. M. R. D. was supported in part by the U.S. Army Research Laboratory and the U.S. Army Research Office under Contract No. W911NF-13-1-0390.

## 5 Appendix

### 5.1 Calculation of Fisher Information, Linear Stage

All variability after the linear stage is Gaussian; thus, the Fisher information can be expressed in the form [1, 26]:

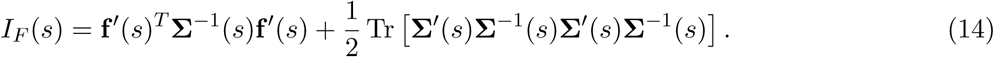

Our immediate goal is to calculate **f** (*s*), the average response of the linear stage, and **Σ**, the covariance between the responses. The output of the *i*th neuron after the linear stage is

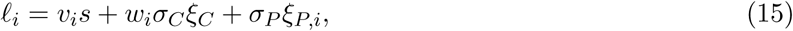

so that the average response as a function of *s* is

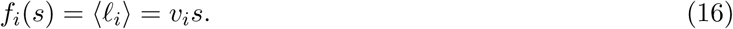

Thus,

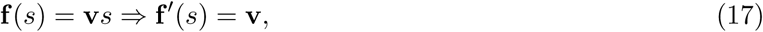

and

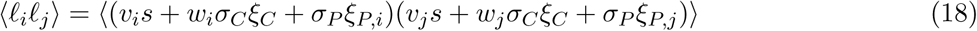

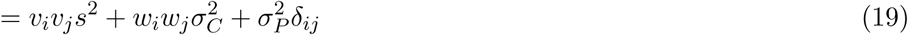

so that

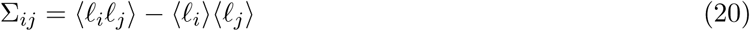

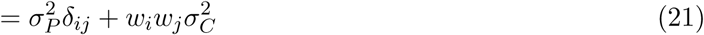

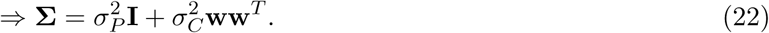

Notice that the covariance matrix does not depend on *s*, so the second term in equation (14) will vanish. We do, however, need the inverse covariance matrix for the first term:

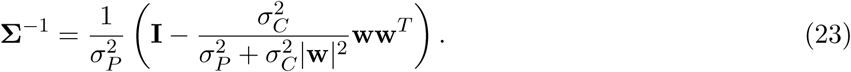

Hence, the Fisher information is

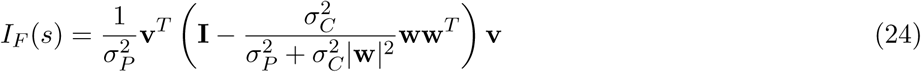

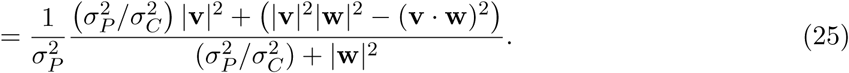

### 5.2 Calculation of Mutual Information, Linear Stage

The mutual information is given by

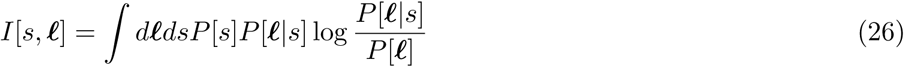

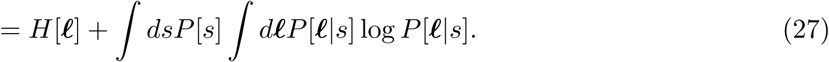

Note that *P* [ℓ] and *P* [ℓ|*s*] are both multivariate Gaussians. The (differential) entropy of a multivariate Gaussian random variable *X* with mean ***µ*** and covariance **Σ** is given by

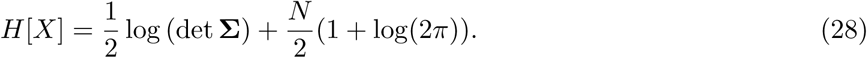

Therefore, by the Gaussianity of the involved distributions,

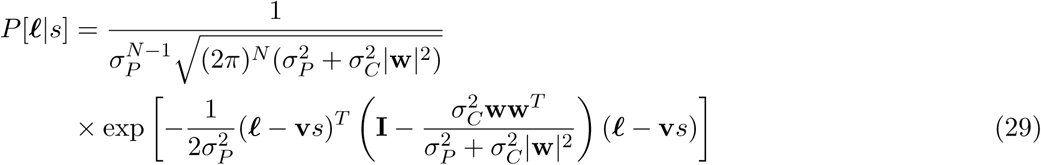

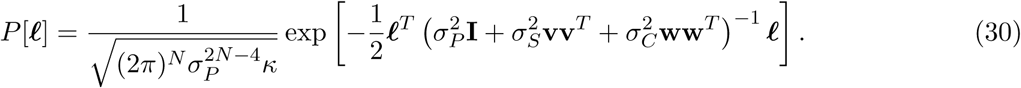

where

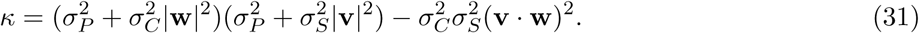

Thus,

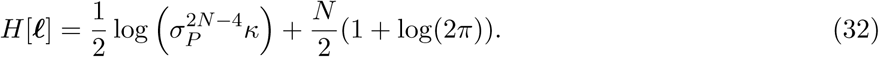

and

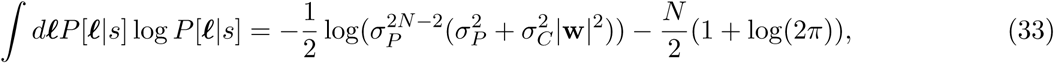

which is notably independent of *s*. Thus, the integral over *s* will marginalize away. We are left with

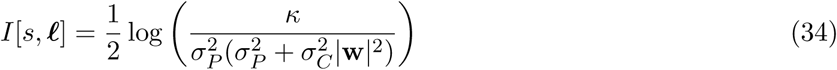

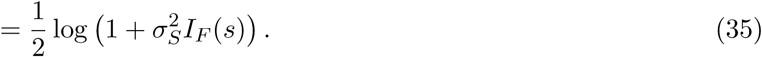

### 5.3 Calculation of Fisher Information, Quadratic Nonlinearity

We repeat the calculation of the first section, but after the nonlinear stage. In this case, we consider a quadratic nonlinearity. Instead of the Fisher information, we calculate the linear Fisher information (since it is analytically tractable). The output of the network is

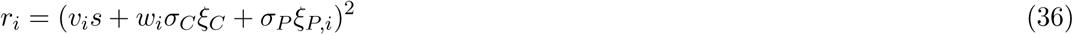

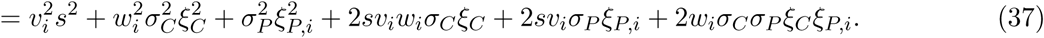

Thus, the average is then

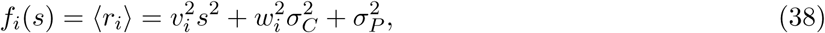

which implies

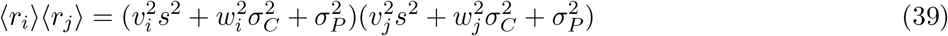

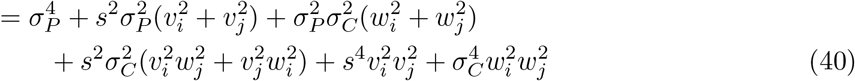

Next, the covariate can be written as

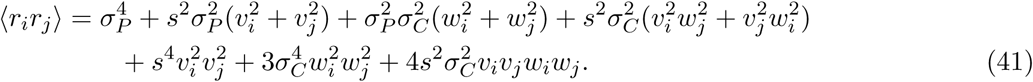

The off diagonal terms of the covariance matrix are then

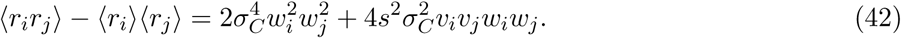

Lastly, the variance of *r*_*i*_ (the diagonal terms of the covariance matrix) is given by

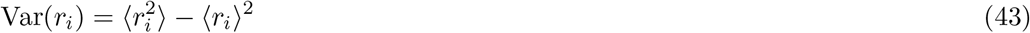

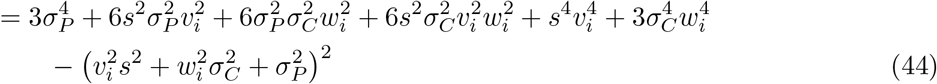

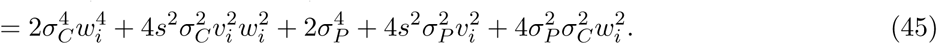

Thus, the total covariance, which takes the variance into consideration, is

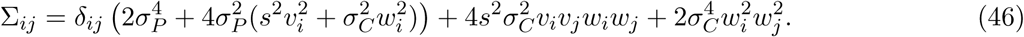

In vector notation, this can be expressed as

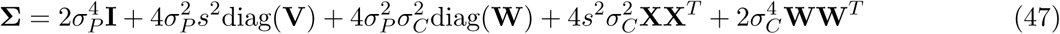

where

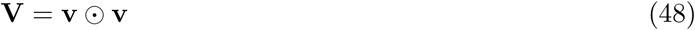

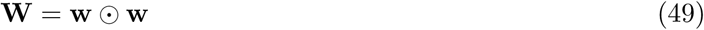

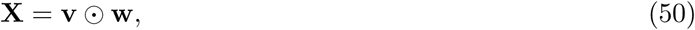

where ⊙ indicates the Hadamard product (element-wise product). We now proceed to the linear Fisher information:

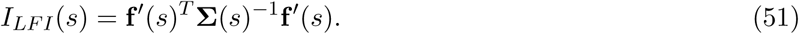

We start by calculating the inverse covariance matrix, which we will achieve with repeated applications of the Sherman-Morrison formula [43]. We can write

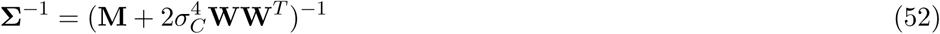

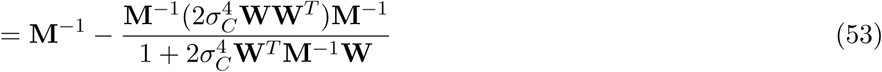

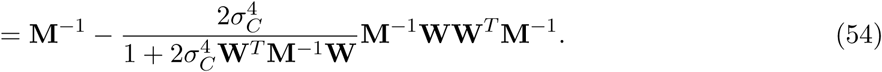

Where

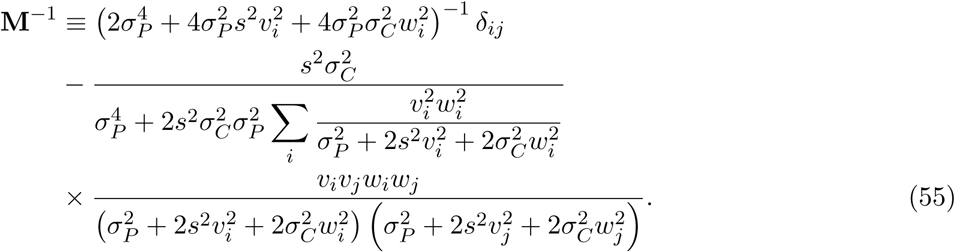

Note that

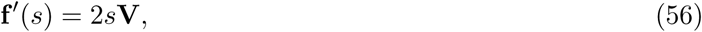

so the Fisher information is

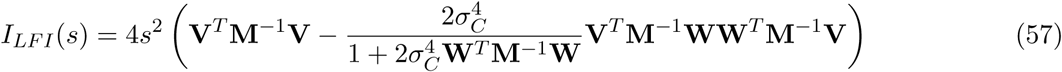

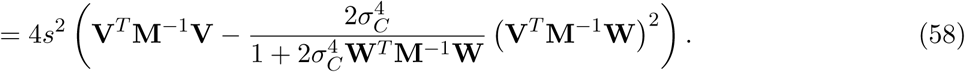

To facilitate the matrix multiplications, we will define the following notation

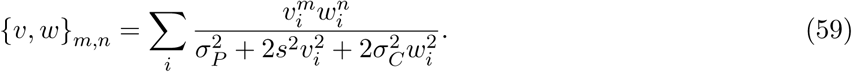

Thus,

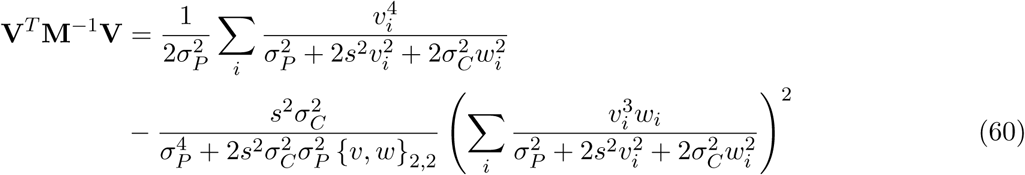

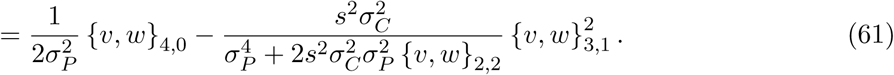

Furthermore,

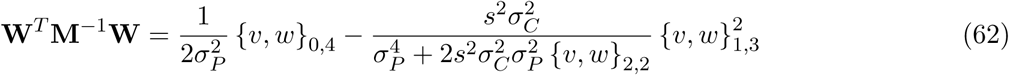

and finally

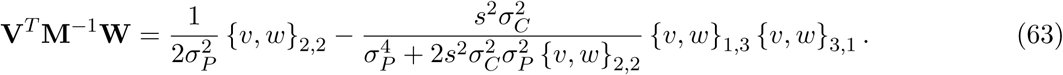

Inserting this expression into equation (58) and simplifying, we can write the Fisher information as

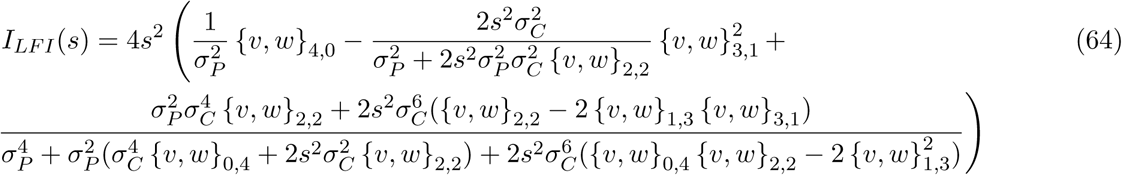

## Footnotes

1 https://github.com/pssachdeva/noise_diversity

